# Behavioural responses to chemical cues of predators differ between fire salamander larvae from two different habitats

**DOI:** 10.1101/2020.10.23.352369

**Authors:** Luca G. Hahn, Pia Oswald, Barbara A. Caspers

## Abstract

Predation is one of the strongest selection pressures, forcing prey organisms to detect predators and to display various antipredator behaviours, such as refuge-use or decreased activity. To recognise predators, chemosensory cues play a pivotal role, particularly in aquatic ecosystems. However, it is less known whether the ability to use these cues to respond with adequate antipredator behaviour varies between individuals occupying different habitats that are dissimilar in predation risk. Using field experiments, we examined antipredator behaviour of larval fire salamanders (*Salamandra salamandra*) from two different habitats, ponds and streams. Among other differences, ponds and streams are inhabited by habitat-specific predators, such as alpine newts (*Ichthyosaura alpestris*) occurring in ponds. We exposed larvae from both habitats to either chemical cues from alpine newts or a blank control (tap water) and investigated potential differences in their behavioural responses in two experiments. Pond larvae, but not stream larvae, became significantly less active when faced with chemical cues from newts compared to those faced with a control stimulus. Moreover, larvae from both habitats tested in water containing chemical cues spent significantly less time outside a shelter than those in control water. Our results demonstrate that larval fire salamanders recognise predatory newts through kairomones and alter their behaviour accordingly. However, experience with predatory newts may not be necessary to differentiate kairomones from control water, but may be beneficial for larvae to further develop their antipredator behaviour, thus representing conformance to a niche.

## Introduction

Predation is a main selection pressure forcing prey organisms to maximise their fitness by recognising and avoiding predators (Lima & Dill, 1990). Consequently, predators can impact behaviour, life history, and morphology of prey individuals and populations (Laforsch & Tollrian, 2004; Lima & Dill, 1990). For example, many prey species from various taxa such as fish, birds, and mammals, form groups to better recognise and escape predators (Krause & Ruxton, 2002). However, the specific mechanisms of how prey organisms use information about predators to perform an antipredator response often remain obscure, limiting our understanding of predator-prey relationships. In these interactions, prey species use their senses to detect a predator, which is crucial to initiate antipredator responses. For example, animals may rely on their visual, auditory, or olfactory system to obtain information about the level of risk (Deecke, Slater, & Ford, 2002; Kelley & Magurran, 2003; Wisenden, 2000). The sensory modalities that prey organisms use to recognise a predator may depend on the habitat. For instance, in aquatic ecosystems, where sight is often restricted, animals may prioritise chemical cues enabling them to exploit information about other species, such as the presence of a potential predator (Kats & Dill, 1998; Kiesecker, Chivers, & Blaustein, 1996). These chemical cues are referred to as “kairomones” (Brown, Eisner, & Whiuaker, 1970), and may stem from the predator itself or from the predator’s diet, potentially conveying information about injured or eaten conspecifics (Chivers & Mirza, 2001; Laurila, Kujasalo, & Ranta, 1997). Using kairomones to avoid predators is widespread across various prey taxa, including fish (Rehnberg & Schreck, 1987), amphibians (Kats, 1988), reptiles (Thoen, Bauwens, & Verheyen, 1986), mammals (Caine & Weldon, 1989), and invertebrates (Bucciarelli & Kats, 2015; Kenison, Weldy, & Williams, 2018; Von Elert & Pohnert, 2000). In case of predator detection, e.g. through kairomones, prey individuals may increase their chances for survival by adjusting their behaviour (Azevedo-Ramos, Sluys, Hero, & Magnusson, 1992; Lawler, 1989). For example, antipredator strategies include decreasing activity, seeking shelter, or escaping (Kavaliers & Choleris, 2001; Wishingrad, Chivers, & Ferrari, 2014). Behaviour could be genetically determined (Bize, Diaz, & Lindström, 2012), learned through experience (Shettleworth, 2010), or most likely, developed by an interaction of both genes and learning (Thornton & Boogert, 2019).

Flexibly learning new behaviours may be costly (Dunlap & Stephens, 2016), but could be beneficial to cope with changing environmental conditions (Sol, Duncan, Blackburn, Cassey, & Lefebvre, 2005). In contrast, more innate behaviour could restrict behavioural flexibility, but could efficiently predispose animals to performing adaptive behaviour (Mery & Burns, 2010). Contentious evidence indicates that antipredator behaviour could be innate, learned, or reliant on both genes and cognition. For instance, amphibian larvae of some species, such as the Mallorcan midwife toad (*Alytes muletensis*), or the common frog (*Rana temporaria*), are able to recognise a predator without prior experience (Griffiths, Schley, Sharp, Dennis, & Román, 1998; Laurila, 2000). Some amphibians, however, such as American bullfrogs (*Lithobates catesbeianus*) or wood frogs (*Lithobates sylvaticus*), can learn to detect and avoid predators (Ferrari & Chivers, 2010; Teixeira & Young, 2014). Therefore, our knowledge regarding the circumstances promoting more innate, learned, or intermediately developed antipredator strategies is contradictory. To fully understand how prey organisms become capable of detecting and avoiding predators both at the proximate and the ultimate scale, experimental research comparing antipredator behaviour of individuals originating from different habitats and experiencing different predatory pressures is crucial.

Different individuals of one species may specialise in and conform to a certain ecological niche, for instance by using a specific habitat type or food source (Bolnick et al., 2003; Dall, Bell, Bolnick, & Ratnieks, 2012). This could minimise intraspecific competition (Polis, 1984). Larval and adult insects and amphibians often use the habitat and food resources differently (Székely, Cogălniceanu, Székely, & Denoël, 2020) and may therefore coexist in higher densities. Partitioning the ecological niche can also occur at finer scales, e.g. within one life history stage (or ‘cohort’) and between individuals (Bolnick et al., 2003; Dall et al., 2012). For instance, individual female fire salamanders deposit their larvae in standing or streaming water bodies within one forest (Caspers, Steinfartz, & Krause, 2015; Steinfartz, Weitere, & Tautz, 2007). As a result, larvae from different habitats are exposed to often markedly dissimilar conditions during their development, potentially affecting their behaviour. In cases where niche partitioning has occurred more recently in the evolutionary history of a species, individuals occupying one niche may not exhibit genetic adaptations to it, making behavioural plasticity fundamental. For instance, individuals from different habitats may respond differently to various stimuli, such as predator kairomones, based on their experience. Several studies investigated the antipredator behaviour of experienced individuals compared to that of naïve individuals (Jackson & Brown, 2011; Mathis, Murray, & Hickman, 2003; Mogali, Saidapur, & Shanbhag, 2012). For example, wild and hatchery-reared juvenile Atlantic salmon (Salmo salar) differed in their antipredator behaviour when tested under seminatural conditions. However, most of these studies were performed with wild-caught and laboratory-raised individuals and do not provide information about differences in anti-predator responses in varying natural habitats. Thus, further research is needed to study how intraspecific niche partitioning affects adaptation and phenotypic plasticity. Moreover, it remains unknown whether individuals of one species populating different habitat types differ in just one specific or in various behaviours.

To investigate how individuals of a species exhibiting niche partitioning differ in their use of predator kairomones and their antipredator behaviour, we conducted field experiments using fire salamander larvae (*Salamandra salamandra*) from two different habitat types, streams and ponds, in the ‘Kottenforst’, a broadleaf forest in Western Germany. Usually, in spring, female fire salamanders deposit their larvae into small first order streams (Thiesmeier, 2004), but in the ‘Kottenforst’ and in other areas, females also choose ephemeral ponds as a larval habitat (Weitere, Tautz, Neumann, & Steinfartz, 2004). The two habitat types differ considerably, thus larvae from both habitats experience different conditions. Predator abundance in streams is relatively low, comprising few dragonfly larvae, whereas predation pressure in ponds is higher due to dragonfly larvae, other invertebrates, and notably, newts (Thiesmeier, 2004). Moreover, ponds are more restricted and unpredictable than streams (Reinhardt, Steinfartz, Paetzold, & Weitere, 2013; Weitere et al., 2004). Streams provide higher food abundance, higher oxygen supply and more stable temperatures (Reinhardt, 2014). The two larval habitats are associated with two genetic clusters (Steinfartz et al., 2007). This phenomenon has not been observed elsewhere in this species and provides an opportunity to study ecological adaptations and niche partitioning in two divergent ecotypes. Salamanders from the two genetic clusters may have already evolved specific behavioural adaptations to their habitat (Weitere et al., 2004). For instance, females from the two clusters differed in their larval deposition behaviour (Caspers et al., 2015) and larvae from the two habitats differed in their risk-taking behaviour (Oswald, Tunnat, Hahn, & Caspers, 2020). Our study system allows to examine if and how individuals from two habitat types behave differently. To investigate whether larvae from ponds and streams differ in their response to predator kairomones, we exposed larvae to chemical cues from predatory alpine newts (*Ichthyosaura alpestris*). In spring, newts temporarily inhabit ponds, but not streams, for reproduction (Joly & Miaud, 1989). Thus, pond larvae are at a higher risk of being preyed on by alpine newts than stream larvae. Kairomone-induced antipredator responses have been investigated in several other salamander species, which often show reduced activity and increased refuge-use (Crane & Mathis, 2011; Kats, 1988; Mathis et al., 2003). However, the potential effect of experience with predators based on the habitat type has rarely been considered in populations where behavioural differences among subpopulations have already been detected. Therefore, our study population provides an opportunity to investigate whether ecotypes differ in various behaviours.

In our study, we intended to quantify whether larvae of the two habitats differ (i) in their activity level and (ii) in their risk-taking behaviour when faced with newt kairomones as these are characteristic antipredator behaviours in larval amphibians (Chivers & Mirza, 2001; Kenison et al., 2018). Pond larvae regularly encounter predatory alpine and other newt species in their habitat (Thiesmeier, 2004), and we hypothesised that chemical cues (kairomones) are a crucial source of information for prey to detect a predator in an aquatic ecosystem, where sight may be unreliable. Consequently, we predicted that in case learning through experience is necessary in fire salamander larvae to recognise the predator through kairomones and to develop adequate antipredator behaviour, pond larvae, but not stream larvae should show antipredator behaviour when being exposed to the chemical cues. More specifically, we predicted that pond larvae, but not necessarily stream larvae, should display reduced activity and be less likely to emerge from a shelter in the treatment containing cues from alpine newts compared to individuals in a tap water control. However, in case of a more innate response, we predicted that larvae from both habitats should exhibit antipredator behaviour by becoming less active and less likely to emerge from a shelter when being faced with kairomones compared to the control.

## Methods

### Data collection and study species

We studied wild fire salamanders in the ‘Kottenforst’, a 30 km^2^ large forest area on an uplifted plateau near Bonn in Western Germany (50°40’ N, 7°7’ E). Fire salamander females deposit up to 70 larvae, sired by up to four males (Caspers et al., 2014) mostly during spring into first order streams or ponds. Larvae stay in their habitat until metamorphosis, which takes 2-3 months under laboratory conditions (Krause, Steinfartz, & Caspers, 2011).From mid-March (18^th^) to mid-May (17^th^) 2019 we collected fire salamander larvae (N = 138) with a dip net from ponds and streams. The sampling locations were two sites of a stream (N = 69; ‘Klufterbach’, 50°41.21’ N, 7°7.60’ E) approximately 150 m apart from each other and four different ponds (N = 69) nearby (about 100 m apart from each other and more than 1 km apart from the stream sites). Larval salamanders from the same water body were kept in large buckets filled with water from their natal habitat for at least one hour after capture to allow them to acclimatise. After the experiments, the larvae were released in their natal water body.

### Behavioural experiments

In two independent two-factorial experiments in the field (each trial 120 s), we tested how larval fire salamanders (N = 138) from two habitats respond to chemical cues (kairomones) from predatory adult alpine newts, living in ponds, as a threat stimulus. Each individual larva was faced with *one* of the two experimental setups (activity test *or* shelter-emergence test). We randomly exposed larvae to a treatment containing chemical stimuli from predatory newts (chemical stimulus) or a blank control (tap water). To obtain the chemical stimuli from the newts, we captured adult alpine newts in ponds nearby and kept five individuals in a jar with 500 ml of tap water for one hour before releasing them. Another jar, filled with the same amount of tap water, served as a blank control. We used tap water in both cases to keep the treatments as neutral as possible, since larvae may respond to multiple chemical cues when using water from the different water bodies. Each day we prepared the water stimuli for the treatments and initiated the experiments immediately afterwards to ensure that cues were present during the tests. Moreover, we renewed the 25 ml of water from the specific treatment for each individual before starting the experiment. The experimenter was blind with respect to the treatment because the two jars containing the water samples (either predator cues or tap water as control) were coded by another person. We observed individual larvae in Petri dishes (9.0 cm diameter) filled with 25 ml water of one of the two treatments. The experiments were performed in a distinct order, by alternating the origin of larvae (pond or stream) and the treatment type (newt water or tap water) to minimise confounding effects. To measure the behaviour and the body length of each salamander larva, we used a stopwatch and millimetre paper (accuracy ± 0.1 mm), respectively. Individuals were transferred from the bucket to the Petri dish with a dip net and acclimatised for one minute before starting the experiment.

#### Activity Test

Amphibian larvae such as wood frog tadpoles became less active when faced with predator cues (Chivers and Mirza 2001). Therefore, we considered activity a suitable proxy for antipredator behaviour and tested for differences between fire salamander larvae from pond and stream habitats. We tested 79 fire salamander larvae (N_pond-control_ = 19, N_pond-newt_ = 21, N_stream-control_ = 20, N_stream-newt_ = 19) by transferring each individual with a dip net into a Petri dish filled with water from either of the two treatments (Figure 1). After releasing larvae into the Petri dish, they were usually active for a few seconds, potentially reflecting a startling response to handling. To avoid a potential influence of this reaction on our test, we waited until the larva was motionless for the first time, usually taking a few seconds. Larvae generally initiated movement soon after handling instead of freezing due to handling. During a trial (120 s) we measured the cumulative time an individual spent moving, considered as activity, by using a stopwatch.

**Figure 1.**
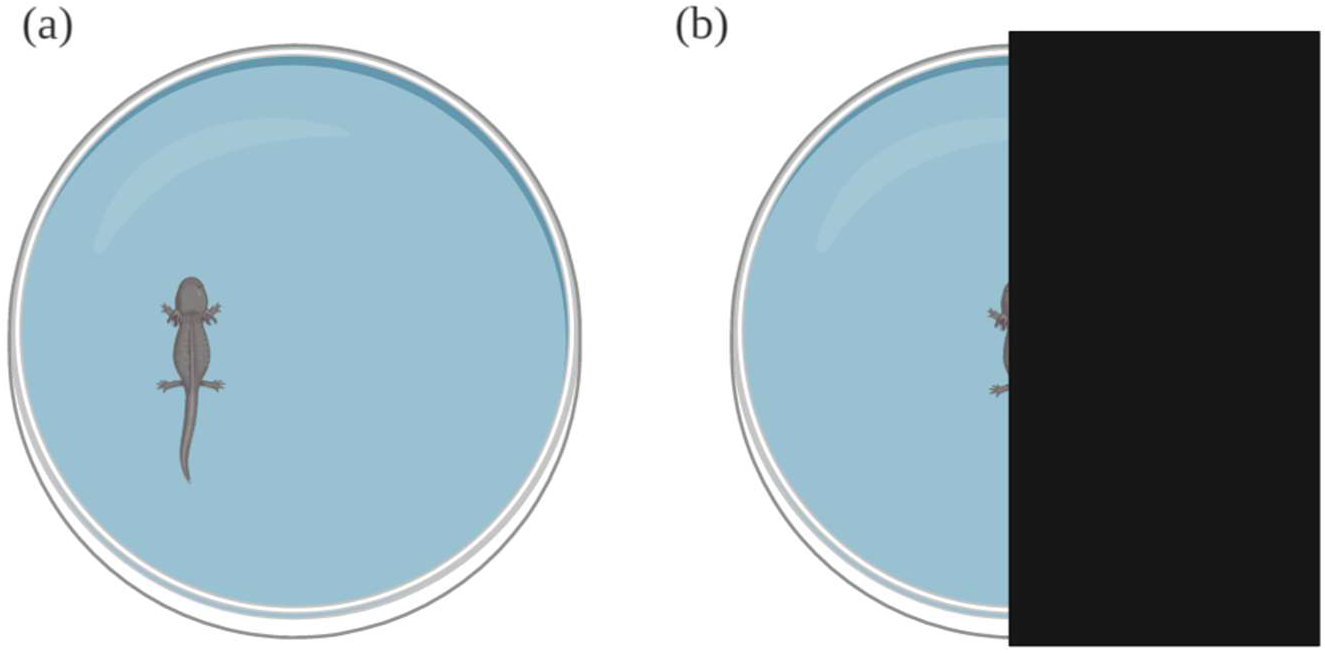
Experimental setup. (a) In the activity test, the cumulative amount of time larvae spent moving was measured. (b) At the start of the shelter emergence test, a larva was located underneath a shelter and we quantified the cumulative amount of time spent outside the shelter. Created with BioRender.

#### Shelter-Emergence Test

For the shelter-emergence test, we placed each larva (N = 59 larvae; N_pond-control_ = 14, N_pond-newt_ = 15, N_stream-control_ = 14, N_stream-newt_ = 16) into a Petri dish containing 25 ml of water. After releasing the larva into the water, we covered one half of the Petri dish with an opaque sheet above the larva to provide a shelter. Thereby, the experiment started with each larva being in the covered area of the Petri dish. During the test (120 s), we measured the time each larva spent outside the shelter with a stopwatch. A larva was considered outside as soon as the head and the forelegs crossed a line on the ground parallel to the shelter. This was considered a higher risk than hiding under the cover.

### Statistical analyses

For all statistical analyses we used the statistical software R (version 4.0.2) (R Core Team, 2020). To investigate possible influences on activity or risk-taking behaviour of fire salamander larvae, we used linear models (LM) with the measured time (s) of the behaviour as a dependent variable. We constructed global models with ‘habitat’ (pond, stream), ‘treatment’ (chemical stimulus or control water) as independent variables and body’ length as a covariate. These models also included an interaction between ‘habitat’ and ‘treatment’ to examine whether the effect of the treatment was dependent on the origin of larvae. Global models were assessed with diagnostic plots to ensure that they met assumptions of linear models, such as normality of residuals, homoscedasticity, and a linear relationship. As the data in the shelter-emergence test violated these assumptions, we used a Box-Cox transformation of the behavioural response variable. After constructing ‘full’ models, we applied step-wise model selection using likelihood ratio tests (LRT) to determine the best model as well as its parameter estimates and significance of predictors. To interpret significant interactions, we used interaction plots and split the dataset into two subsets based on the habitat (one subset per habitat). This allowed us to test separately whether the treatment affected the behaviour of pond and stream larvae. We further compared the body length of larvae from the two habitats by performing a LM to examine whether body length as a dependent variable was different across treatment groups by including an interaction between the two fixed effects ‘habitat’ and ‘treatment’. We calculated means, standard deviations (SD), and standard errors (SE) of the measured variables.

## Results

To test for potential differences in their behavioural response towards chemical cues of a predator, we tested 138 larvae (N_pond_ = 69, N_stream_ = 69) in *one* of two experiments, the activity test *or* the shelter-emergence test. Therefore, larvae were faced with one of two different water treatments, (i) control water, or (ii) chemical stimulus water (i.e. water that had contained a newt).

### Activity Test

We tested 79 salamander larvae (N_pond_ = 40, N_stream_ = 39) with a total body length ranging from 2.65 to 4.35 cm for their activity. Larvae from the two habitats differed significantly regarding their body length (LM, β ± SE(β) = −0.225 ± 0.093, F_1,77_ = 5.881, *P* = 0.018). Salamanders from ponds (mean = 3.67 cm, SD ± 0.47 cm, SE ± 0.07 cm) were significantly larger than those from the streams (3.44 cm, SD ± 0.35 cm, SE ± 0.06 cm). An interaction between the larval habitat and the treatment significantly affected activity levels of fire salamander larvae (LM, β ± SE(β) = 15.967 ± 4.474, F_1,74_ = 12.738, *P* < 0.001; Table 1; Figure 2). Pond larvae differed significantly in their activity depending on the treatment, i.e. pond larvae in the newt treatment spent less time moving than those in the blank control (LM, β ± SE(β) = −18.687 ± 3.447, F_1,37_ = 29.383, *P* < 0.001; revealed by splitting the dataset based on the two habitat groups). In contrast, no such difference was found between the two treatment groups of stream larvae (LM, β ± SE(β) = −3.251 ± 2.820, F_1,36_ = 1.330, *P* = 0.257). Larval salamanders from the ponds were more active than those from the streams (Table 1; Figure 2). Pond larvae faced with the control treatment were the most active ones (34.81 s, SD ± 12.79 s, SE ± 2.93 s) and were more than twice as active as those in the newt treatment (14.57 s, SD ± 11.66 s, SE ± 2.55 s). Stream salamanders in the control treatment spent 6.94 s active (SD ± 10.35, SE ± 2.31 s), those in the newt treatment moved on average 3.51 s (SD ± 6.86 s, SE ± 1.57 s). Activity of the larvae was positively associated with their body length, with larger individuals being more active (LM, β ± SE(β) = 9.892 ± 2.757, F_1,74_ = 12.873, *P* < 0.001, Table 1).

**Figure 2.**
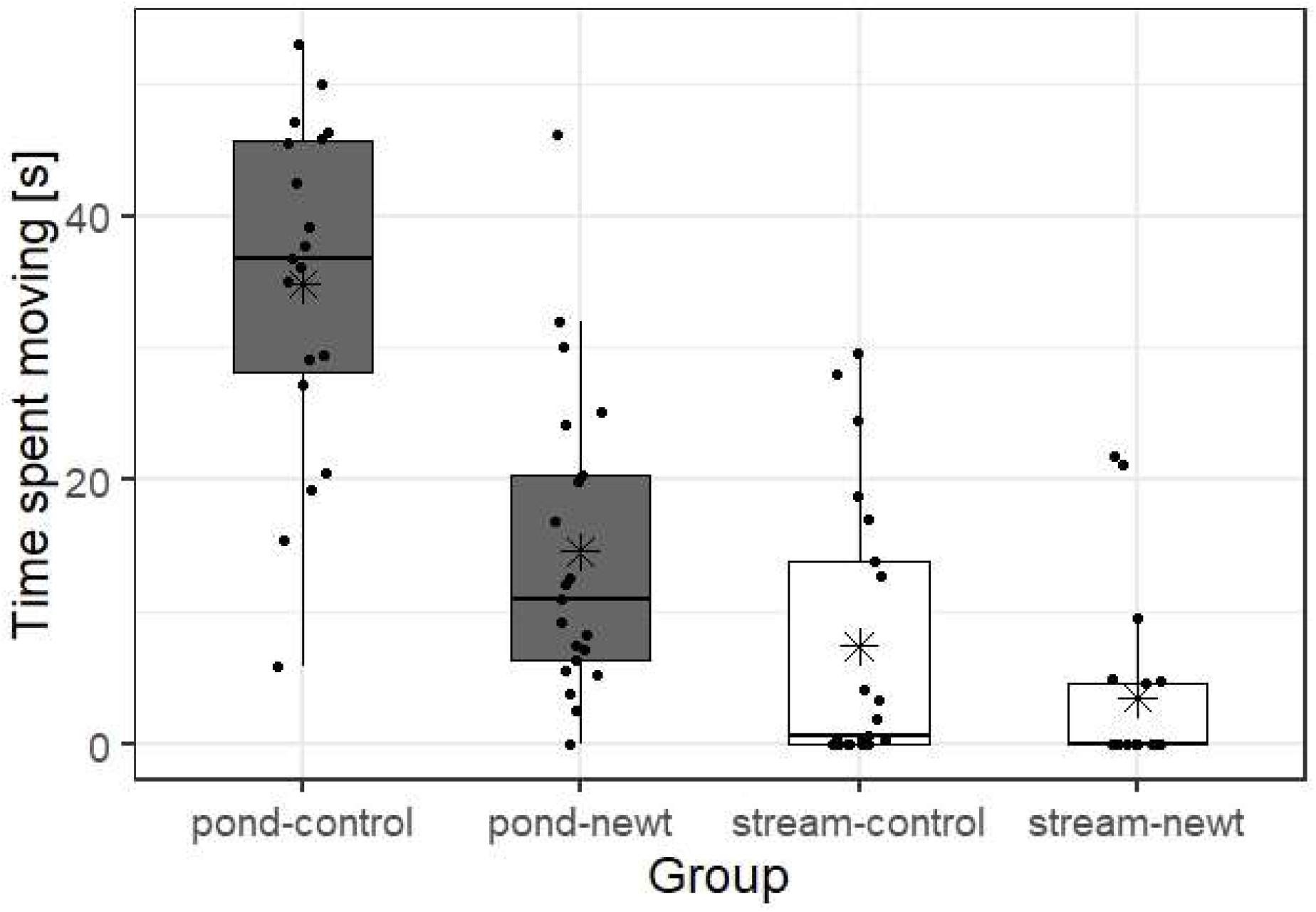
Activity of larval fire salamanders from the four treatment groups (N_pond-control_ = 19, N_pond-newt_ = 21, N_stream-control_ = 20, N_stream-newt_ = 19). Each larva was faced with either the treatment containing chemical stimuli from Alpine newts (N = 40) or the control treatment (N = 39). There was a significant interaction between habitat and treatment. Horizontal lines indicate the median, asterisks the mean. The edges of the boxes limit the first and the third quartile. One dot represents one individual.

### Shelter-Emergence Test

In the shelter-emergence experiment we tested 59 larvae (N_pond_ = 29, N_stream_ = 30) with a total body length ranging from 2.45 to 4.70 cm. Again, in this experiment, pond larvae (3.50 cm, SD ± 0.46 cm, SE ± 0.08 cm) were significantly larger than stream larvae (3.23 cm, SD ± 0.43 cm, SE ± 0.08 cm) (LM, β ± SE(β) = −0.275 ± 0.115, F_2,57_ = 5.730, *P* = 0.020). The larval habitat (LM, β ± SE(β) = 82.92 ± 37.03, F_4,55_ = 6.089, *P* = 0.017; Table 2) and the treatment (LM, β ± SE(β) = 73.18 ± 36.38, F_4,55_ = 4.792, *P* = 0.033, Table 2) but not their interaction (LM, β ± SE(β) = −35.03 ± 51.00, F_4,55_ = 0.472, *P* = 0.495), significantly affected the behaviour of fire salamander larvae (Figure 3; please note that the behavioural variable for this test was Box Cox transformed). In general, pond larvae spent more time outside the shelter (30.87 s, SD ± 27.12 s, SE ± 5.04 s) than stream larvae (9.8 s, SD ± 25.64 s, SE 4.68 ± s). Salamanders from both habitats exposed to the chemical stimuli from newts spent less time outside the shelter than those in the control. More specifically, pond larvae in the control stayed 32.9 percent of the time (39.5 s, SD ± 29.32 s, SE ± 7.84 s) in the uncovered area. Pond salamanders in the newt treatment were visible in the open sector for 19.0 percent of the time (22.81 s, SD ± 22.99 s, SE ± 5.94 s). Stream salamanders in the control treatment spent 15.46 s (SD ± 34.32 s, SE ± 9.15 s, 12.9 %) outside the shelter, whilst those in the newt treatment did so for 4.85 s (SD ± 14.11 s, SE ± 3.53 s, 4.0 %). There was a non-significant trend regarding body length, with larger larvae spending more time in the open area (LM, β ± SE(β) = −56.77 ± 29.41, F_4,55_ = 3.502, *P* = 0.067).

**Figure 3.**
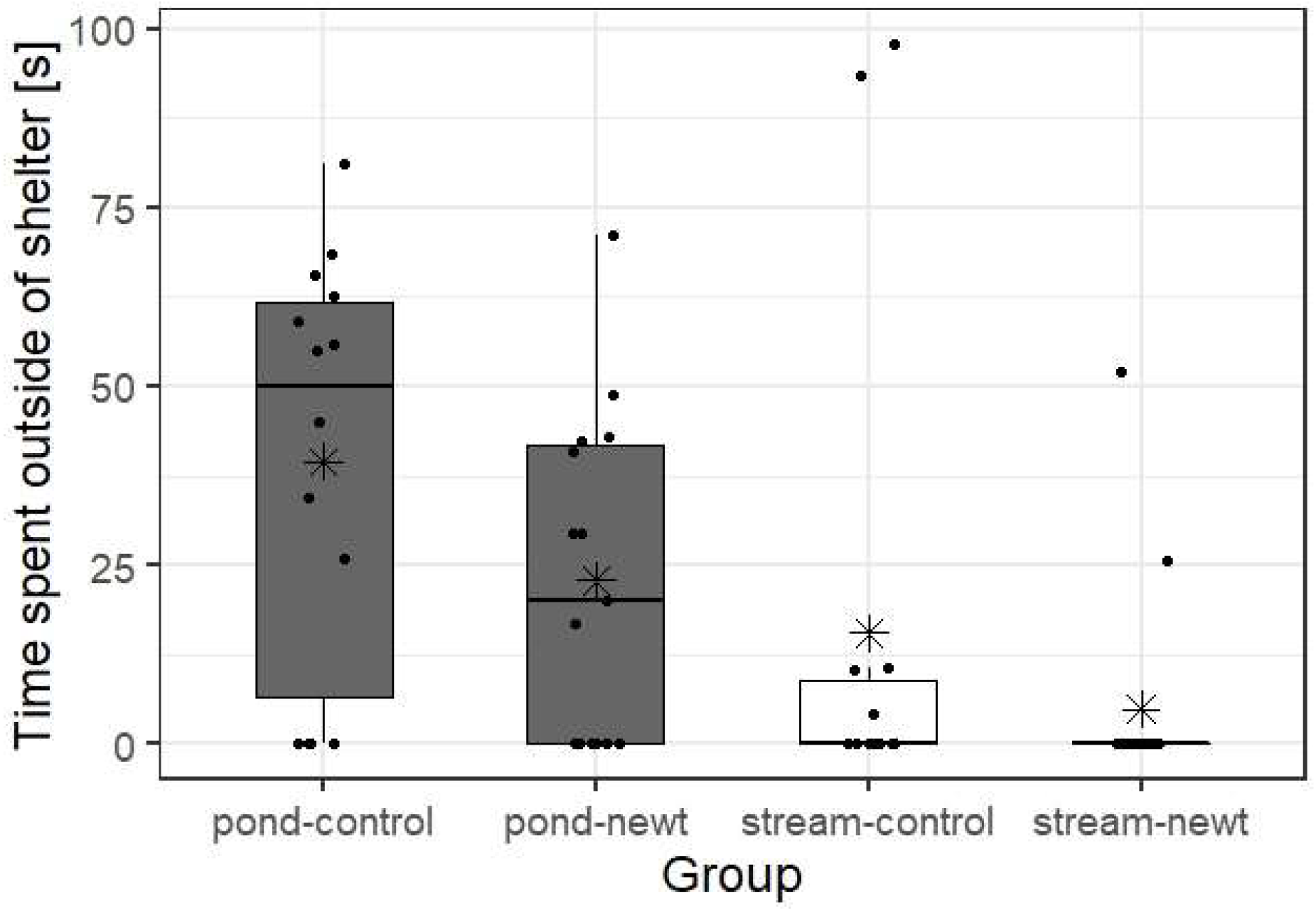
Time salamander larvae from the four experimental groups (N_pond-control_ = 14, N_pond-newt_ = 15, N_stream-control_ = 14, N_stream-newt_ = 16) spent outside the shelter in a trial lasting 120 s. There was a significant impact of treatment and habitat treatment, but not of the interaction of habitat * treatment. Each dot represents one individual. The horizontal lines in the boxes stand for the median, the asterisks indicate the mean. The boxes limit the first and the third quartile.

## Discussion

This study revealed significant differences in the behaviour towards chemical cues from alpine newts between fire salamander larvae originating from two ecological different habitats, ponds and streams. Larvae from ponds, but not from streams, were significantly less active when confronted with chemical cues from newts compared to those individuals encountering a control stimulus. However, in the shelter emergence test, larvae from both habitats showed less risk-taking behaviour in the newt treatment than those in the control treatment. Therefore, this study demonstrates larval fire salamanders from different habitats use chemical cues (i.e. kairomones) to detect and avoid predatory newts, but they do so differently, potentially based on different experiences with this predator due to their natal habitat.

As predicted, larval fire salamanders responded to information conveyed by predator chemical cues with antipredator behaviour. Predators impose a strong selection pressure on prey organisms, forcing them to evolve and develop appropriate antipredator strategies including behaviour, life-history, and morphology (Lima & Dill, 1990). Particularly through behaviour, prey individuals can avoid predators relatively flexibly and therefore maximise their fitness (Curio, 1976). To perform an adequate antipredator behaviour, however, prey animals have to recognise the predator first. Prey species have evolved senses to inform themselves about the level of risk (Deecke et al., 2002; Kelley & Magurran, 2003; Wisenden, 2000). To assess predation risk in aquatic ecosystems, where sight and acoustics may be impaired, prey species may prioritise chemical cues instead (Brown et al., 1970; Hickman, Stone, & Mathis, 2004). Accordingly, fire salamander larvae were able to adjust their behaviour in the presence of kairomones without de facto encountering newts. These findings are congruent with results from previous studies on antipredator behaviour in prey exposed to predator kairomones (REF). A widespread antipredator response to chemical cues in various taxa is to become less active to minimise attracting the predator’s attention (Azevedo-Ramos et al., 1992; Lawler, 1989). For instance, larval ringed salamanders (*Ambystoma annulatum*) were less active in field and laboratory assays when faced with kairomones from predatory eastern newts (*Notophthalmus viridescens*) (Mathis et al., 2003). Similarly, common frog tadpoles also reduced their activity after exposure to kairomones from perch (*Perca fluviatilis*) and dragonfly larvae (*Aeshna juncea*) (Laurila 2000). In addition to reduced activity, another common behaviour to avoid predators is seeking a refuge (Kats, 1988; Van Buskirk & Schmidt, 2000). When subjected to caged predatory dragonfly larvae, larval newts (*Triturus* sp.) for example, hid more often (Van Buskirk & Schmidt, 2000). Equally, small mouthed salamander larvae (*Ambystoma texanum*) sought shelter when confronted with chemical cues from predatory green sunfish (*Lepomys cyanellus*) (Kats, 1988). As expected, the shelter-emergence test in this study revealed fire salamander larvae were less likely to emerge from the shelter when exposed to newt stimuli. We observed a change in risk-taking behaviour (shelter emergence) in larvae of both habitat types, indicating a innate ability to recognise chemical cues from newts. In addition we found reduced activity and risk-taking behaviour particularly pronounced in larval fire salamanders from ponds.

Predation pressure differs between the two habitats, and newts only occupy ponds, which might consequently result in differences in experience. Thus, differences in experience may thus explain why pond larvae became less active in the newt treatment, whereas stream larvae did not. These findings could support our hypothesis that learning through experience may be necessary in larval fire salamanders to glean information about the presence of a predator and to act upon this information. Pond larvae, but not stream larvae, may have learned that reducing activity is beneficial to avoid predatory newts. However, larvae from both habitats seemed to be able to recognise the chemical cues, as all larvae, irrespective of their habitat, spent less time outside the shelter when exposed to chemical cues. We therefore conclude that larvae may innately be capable of detecting chemical cues and acting accordingly, but additional experience and learning may be necessary for more sophisticated antipredator behaviour to develop. Accordingly, current evidence suggests antipredator strategies in larval amphibians (and other taxa) may be underpinned by both genes and learning. On the one hand, some studies indicate predator recognition is innate in amphibian larvae. For instance, naïve tadpoles of the Mallorcan midwife toad became less active when exposed to stimuli from viperine water snakes (*Natrix maura*) (Griffiths et al., 1998). Additionally, naïve tadpoles of the common frog were less active when encountering chemical cues from two predators (Laurila, 2000). By contrast, other studies demonstrate amphibian larvae can learn to recognise and avoid predators. For instance, larvae of different amphibian species learned to respond to chemical cues from different predators (Murray, Roth, & Wirsing, 2004). Another example are captively bred American bullfrog tadpoles, which learned to avoid a model avian predator (Teixeira & Young, 2014). Moreover, wood frog embryos experiencing kairomones displayed antipredator behaviour later on during the larval stage (Ferrari & Chivers, 2010). Promoting a more comprehensive view, other studies (including this one) state recognising predators may be driven by both genetic and learning mechanisms (Crane & Mathis, 2011; Epp & Gabor, 2008). For instance, both mechanisms seemed relevant in hellbender salamanders (*Cryptobranchus alleganiensis*) (Crane & Mathis, 2011). In this species, individuals were able to recognise natural predators without having encountered them, and they additionally learned to avoid unknown predators through training. In addition, a similar conclusion was drawn in a study on the San Marcos salamander (*Eurycea nana*) (Epp & Gabor, 2008). Congruent with these findings, this study indicates both innate predator recognition and learning through experience could be required for antipredator behaviour in larval fire salamanders. We acknowledge that the predictions made earlier are somewhat simplistic and may neglect a sophisticated phenomenon, whereby antipredator behaviour could be underpinned by both genetics and cognition.

Differences between pond and stream larvae might alternatively be explained by genetic adaptations to the two habitats rather than learning. For example, the water current is different between ponds and streams and this may influence activity levels of larvae in general. Indeed, we found that stream larvae were generally less active than pond larvae. Lower amounts of activity in streams might be an adaptation to avoid drift. Similarly, reduced activity levels in pond larvae in the presence of newt stimuli could reflect a genetic adaptation instead of learning. These alternative explanations could be in accordance with the two genetic clusters corresponding to the two different larval habitats in our study population (Steinfartz et al. 2007, Hendrix et al. 2017). Thus, differences in behaviour between larvae from both habitats might be explained by genetic differences, but this is pure speculative at the moment as we did not take genetic samples. The latter could be tested by using naïve larvae from both habitats that were raised in a laboratory under the exact same conditions without predator experience.

Another variable influcening larvae’s behaviour was their body length, with larger individuals being more active and tending more to leave the shelter. Smaller individuals could be at greater predation risk; reduced activity and shelter-emergence behaviour may minimise this risk. Moreover, larger individuals may have to take more risks to meet their energetic needs. In this study, pond salamander larvae were larger than stream larvae, which could have confounded the effect of the treatment on the behaviour. However, the treatment effect remained despite the effect of body length, a covariate in our models. Previous studies have demonstrated a similar effect of body size on behaviour (Eklöv, 2000; E. T. Krause et al., 2011; Puttlitz, Chivers, Kiesecker, & Blaustein, 1999). Nevertheless, in this study, the treatment (i.e. newt stimuli) impacted the behaviour significantly beyond any size effect.

## Conclusions

Predator-prey interactions are omnipresent in various ecosystems, pressuring prey species to exhibit a plethora of antipredator strategies to recognise and avoid predators. For instance, prey are able to regulate their behaviour in the presence of chemical cues predators release and can use this information to modulate their individual behaviour (Müller, Caspers, Gadau, & Kaiser, 2020). Our results suggest infochemicals may be a vital source of information for prey, such as larval salamanders, that may be innately predisposed to recognise risk imposed by predator cues. Moreover, larval salamanders may be capable of learning to further develop the response because only pond larvae, most likely experienced with chemical cues form newts, but not pond larvae became less active in the newt treatment. Furthermore, this study highlights that ecological niche partitioning and conformance of individuals to their niche can correspond to differences in behaviour.

## Acknowledgements

We thank Benjamin Tunnat for assistance in the field.

## Author contributions

All authors contributed to study design and conception. Data collection in the field was performed by L. G. H. and P. O. Data analysis and the first draft of the manuscript were undertaken by L. G. H. All authors commented on previous versions of the manuscript and approved final submission.

## Availability of data and code

Raw data and code used are made publicly available.

## Ethics approval

We thank the State Agency for Nature, Environment and Consumer Protection (LANUV; reference number 2019: 81-02.04.2019.A112), the nature reserve authority of the Stadt Bonn and the forest warden’s office in Bonn for the permissions to run our experiments in the Kottenforst. Experiments comply with the current laws of Germany. After the experiment, the larvae and newts were released at the site of capture, i.e.in the same water body.

## Consent to participate

Not applicable.

## Consent for publication

Not applicable.

